## Supplementary Figures for "Differential RNA editing between epithelial and mesenchymal tumors impacts mRNA abundance in immune response pathways"

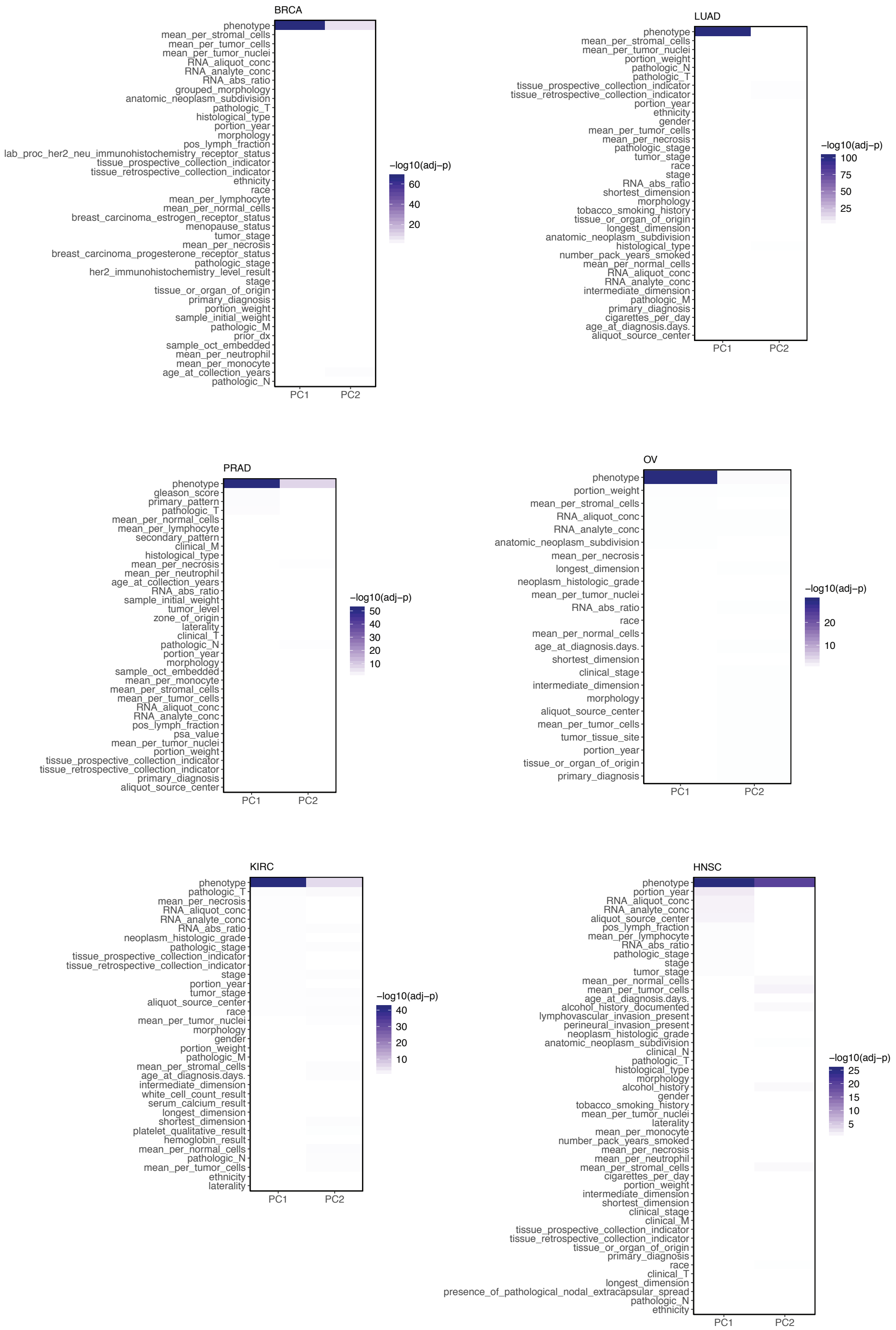


**Supplementary Fig. 1 Differential Editing Not Confounded by Metadata.** Heatmaps of significance (log10-transformed adjusted p-values) of correlations between the top two principal components and E/M phenotype among metadata fields in each cancer type. Darker color indicates smaller p-value and stronger association.

A

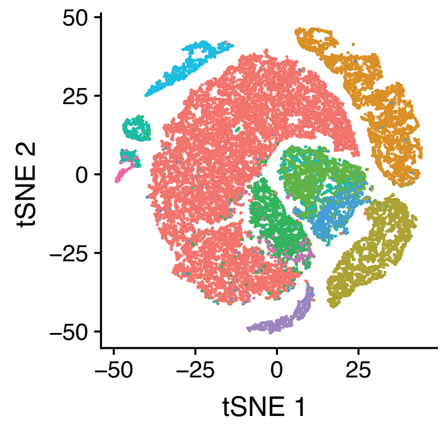


| Cluster | Matching cell types | Matching markers |
|---|---|---|
| | T cell | CD3D |
| | Myeloid | LYZ,MARCO,CD68,FCGR3A |
| | Cancer | EPCAM |
| | Cancer | EPCAM |
| | B cell | CD79A |
| | Cancer | EPCAM |
| | Fibroblast | COL1A1,DCN,COL1A2,C1R |
| | B cell | CD79A |
| | Endothelial Cell | CLDN5,FLT1,CDH5,RAMP2 |
| | Mast | TPSAB1,CPA3,HPGDS,KIT |
| | Alveolar; Epi; Cancer | FOLR1;CAPS,TMEM190,PIFO,SNTN;EPCAM |

B

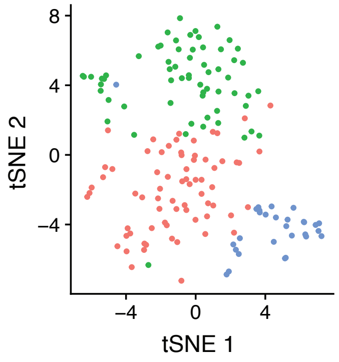


| Cluster | Matching cell types | Matching markers |
|---|---|---|
| | NA | NA |
| | Epithelial | CAPS,TMEM190,SNTN |
| | Alveolar | SFTPB |

C

| Cell Type | Count |
|---|---|
| T cell | 16354 |
| Cancer | 6526 |
| Myeloid | 4063 |
| B cell | 2770 |
| Fibroblast | 998 |
| Endothelial Cell | 771 |
| Mast | 285 |
| Epithelial | 62 |
| Alveolar | 29 |

D

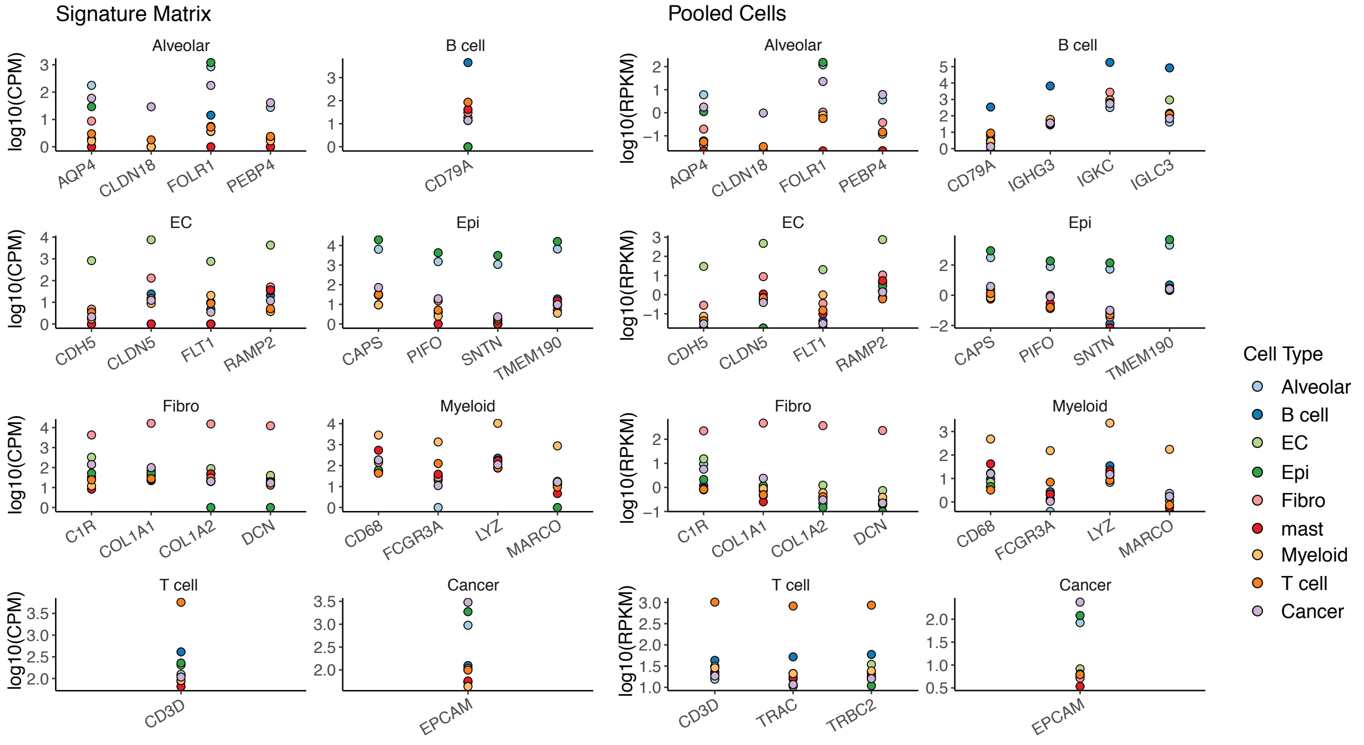


**Supplementary Fig. 2 Clustering of single cells from three lung cancer tumors. A.** TSNE projection of cells based on expression profiles, with color indicating cluster identity (left). Cell types were assigned to clusters by matching differentially expressed genes of clusters to known cell type markers (right). **B.** TSNE projection of only cells from cluster 10 to further refine cell type assignment (left). Similar to **A**, cell types were labeled using differentially expressed genes that matched cell type markers (right). **C.** Counts of cells for each cell type after 2 rounds of clustering and cell type assignment (**A** and **B**). **D.** Log2-transformed expression values of marker genes across cell types. Signature matrix on the left indicates expression values assigned for each cell type by CIBERSORTx. On the right, Pooled Cells indicate that expression values were calculated from pooling reads from cells of the same type together.

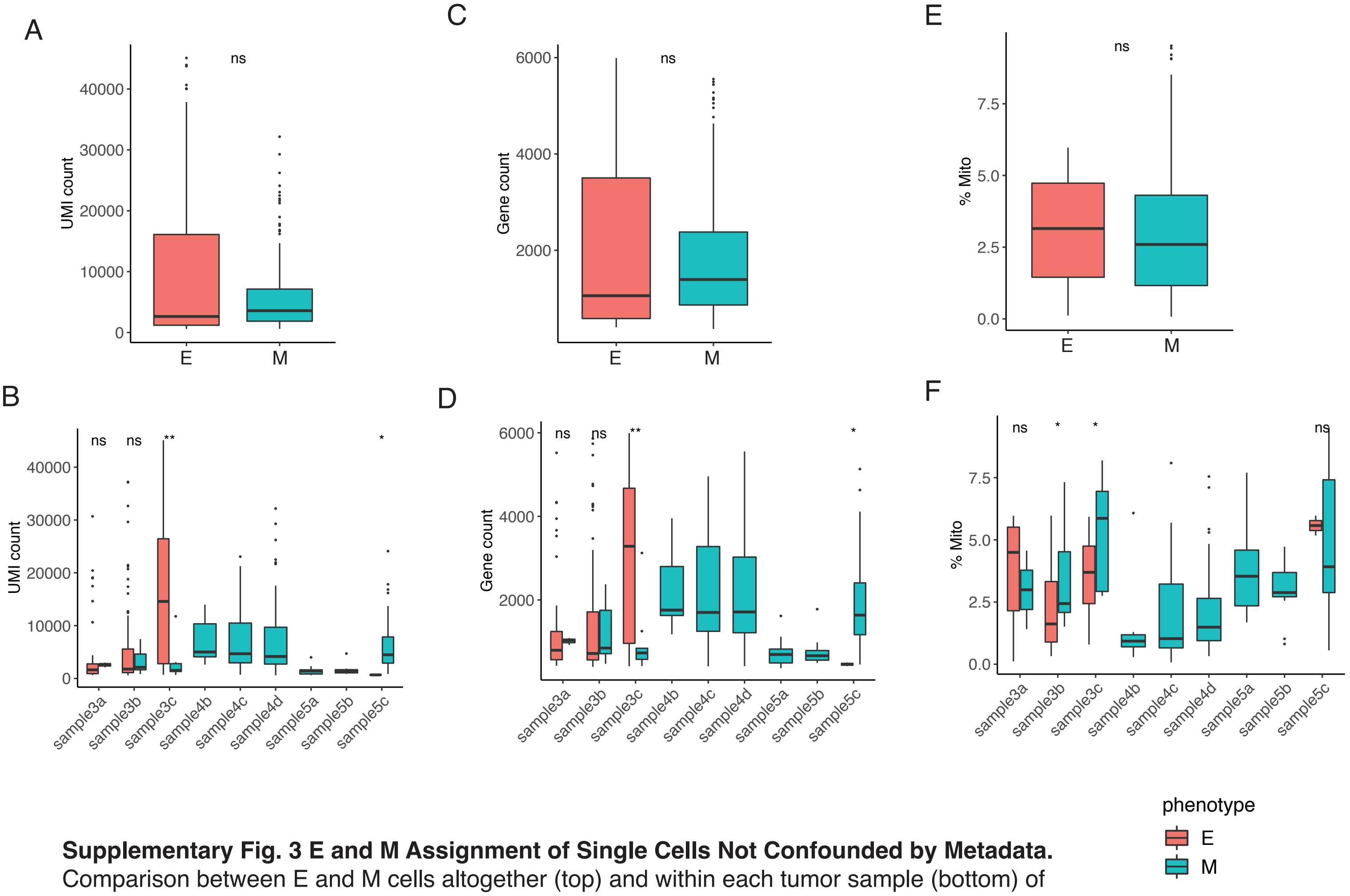


**Supplementary Fig. 3 E and M Assignment of Single Cells Not Confounded by Metadata.** Comparison between E and M cells altogether (top) and within each tumor sample (bottom) of metadata fields: UMI count (**A**-**B**), gene count (**C**-**D**), and percent of reads mapping to the mitochondrial genome (**E**-**F**). Metadata values were compared by Mann Whitney U tests, and significance of p-values are shown. ns: $p > 0.05$, * $p \leq 0.05$, ** $p \leq 0.01$.

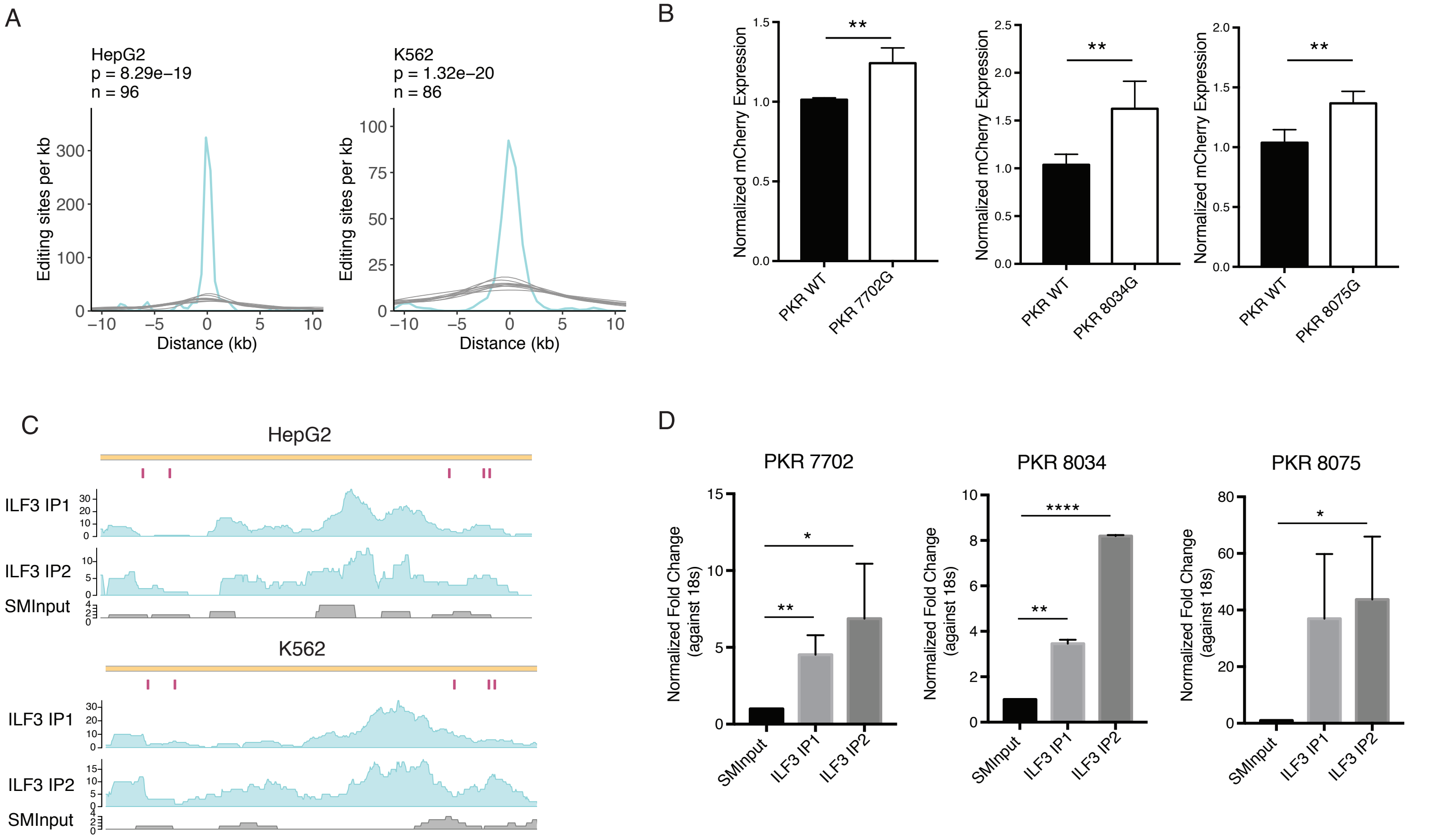


**Supplementary Fig. 4 ILF3 Regulates mRNA Abundance through RNA Editing. A.** Histogram of distances between differential editing sites in editing-correlated genes and the closest ILF3 eCLIP peaks in HepG2 and K562 cells (turquoise), up to 10 kb. Gray curves represent distances for 10 sets of randomly picked A's in the same genes as differential editing sites. Number of differential editing sites is given by n for each cell line. P-value was calculated by comparing the area under the curve (AUC) of the distance distribution for differential editing sites to a normal distribution fit to the AUC values of 10,000 sets of random gene-matched A's. **B.** Normalized mCherry expression for nonedited or edited versions of sites in the 3'UTR of PKR in A549 cells. Normalized expression values were compared between edited and nonedited versions by two-sided t-test. **$p<0.01$. **C.** Read coverage of ILF3 eCLIP-seq in HepG2 and K562 cells for two biological replicates (ILF3 IP1 and ILF3 IP2, turquoise) and size-matched input (SMInput, gray) in each cell line. The five validated 3' UTR editing sites affecting PKR mRNA abundance in A549 cells are labeled in magenta. **D.** Validation of PKR eCLIP signal overlapping three editing sites. PKR expression was measured by qRT-PCR in the IP or SMInput samples and normalized against the expression of 18s rRNA. Three technical replicates were performed (other than two replicates for 8034). P-value calculated by t-test. *$p<0.05$, **$p<0.01$, ****$p<0.0001$.
